# Identifying conversion efficiency as a key mechanism underlying food webs evolution: A step forward, or backward?

**DOI:** 10.1101/640433

**Authors:** Coralie Fritsch, Sylvain Billiard, Nicolas Champagnat

**Affiliations:** Université de Lorraine, CNRS, Inria, IECL, UMR 7502, F-54000 Nancy, France; Université de Lille, CNRS, UMR 8198, Evo-Eco-Paleo, F-59655 Villeneuve d’Ascq, France

**Keywords:** food webs models, trophic interactions, networks, community ecology, ecosystem, adaptive dynamics

## Abstract

Body size or mass is generally seen as one of the main factors which structure food webs. A large number of evolutionary models have shown that indeed, the evolution of body size (or mass) can give rise to hierarchically organized trophic levels with complex between and within trophic interactions. However, because these models have often very different assumptions, sometimes arbitrary, it is difficult to evaluate what are the real key factors that determine food webs evolution, and whether these models’ results are robust or not. In this paper, we first review the different adaptive dynamics models, especially highlighting when their assumptions strongly differ. Second, we propose a general model which encompasses all previous models. We show that our model recovers all previous models’ results under identical assumptions. However, most importantly, we also show that, when relaxing some of their hypotheses, previous models give rise to degenerate food webs. Third, we show that the assumptions made regarding the form of biomass conversion efficiency are key for food webs evolution, a parameter which was neglected in previous models. We conclude by discussing the implication of biomass conversion efficiency, and by questioning the relevance of such models to study the evolution of food webs.

## 1 Introduction

One of the goals of theoretical ecology is to decipher the mechanisms underlying the emergence, topology and stability of trophic food webs. In a context of global change, it is especially important to develop models predicting the evolution of ecological systems in response to temperature change, harvesting increase or populations fragmentations through for instance the evolution of individuals’ traits such as size and feeding rates (Brown et al., 2004; Woodward et al., 2005). A large variety of models have been developed to address this question (reviewed by Brännström et al., 2011).

One class of models is composed of variations of the seminal work by Loeuille and Loreau (2005) (see Section 2 and Table 1 for a compilation). These models study the evolution of a food web, where species are structured by a finite number of continuous traits, including body size or body mass and predation preferences. Population dynamics follows Lotka-Volterra models where three types of interactions are generally considered, within and/or between species: competition for resources, competition independent of the resources, and predation. These models are inspired by the adaptive dynamics framework (Metz et al., 1996; Geritz et al., 1998), i.e. mutations affecting one or several traits (especially body size or mass) are recurrently introduced into the community. The evolution of traits induced by recurrent mutations and ecological interactions can lead to evolutionary branching. Typically, those models show that a food web can emerge with an increase in species number and a given topology for trophic interactions: a given species can preferentially consume a subset of the extant species and/or the resources.

**Table 1:**
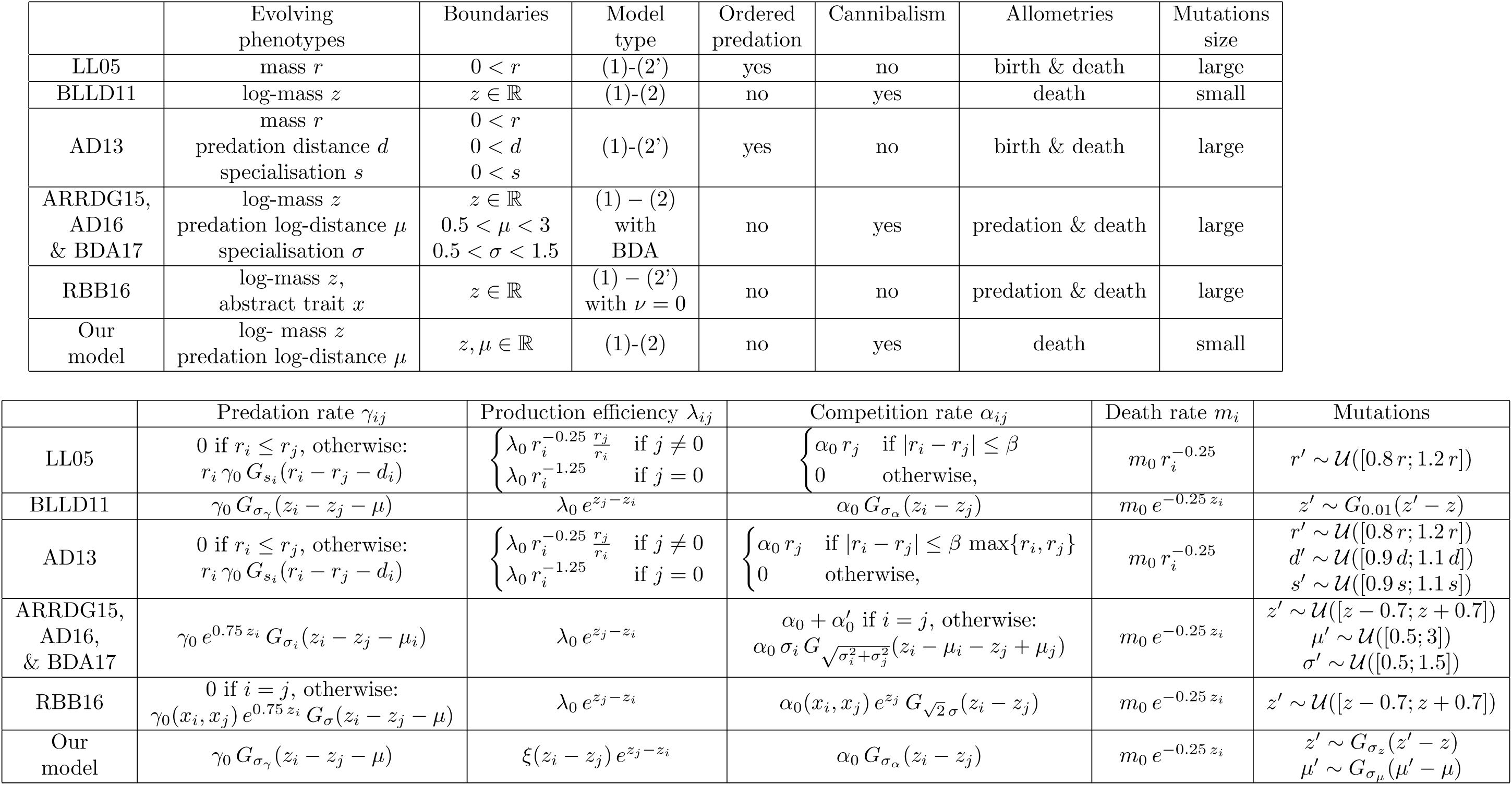
Food web models and predation, production efficiency (biomass conversion + reproduction), competition, death and mutation functions in models of Loeuille and Loreau (2005) (LL05), Brännström et al. (2011) (BLLD11), Allhoff and Drossel (2013) (AD13), Allhoff et al. (2015) (ARRDG15), Allhoff and Drossel (2016) (AD16), Bolchoun et al. (2017) (BDA17), Ritterskamp et al. (2016) (RBB16) and our model (function *ξ* is introduced in Section 5.1). BDA means Beddington-DeAngelis functional response, 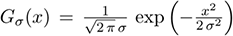 is the centred gaussian density with variance *σ*^2^ and 𝒰 ([*a, b*]) is a uniform distribution on the interval [*a, b*]. When models are expressed in terms of biomass, we convert the differents parameters using (3).

These models have been succesful in showing that 1) the evolution of body size or body mass could be a major mechanism explaining the structure of food webs, and especially the emergence of trophic levels where large species tend to eat small species (e.g. Loeuille and Loreau, 2005; Brännström et al., 2011); 2) diversification in trophic networks are promoted by the number of evolving traits such that predation preference and specialization (Allhoff and Drossel, 2013; Allhoff et al., 2015; Allhoff and Drossel, 2016; Bolchoun et al., 2017) or abstract traits (Ritterskamp et al., 2016); 3) there can exist a turnover of species in trophic networks, with species going extinct and replaced either by new species appeared by mutations, or because of the evolution of the niche of an extant species (Allhoff et al., 2015).

Despite these models are derived from the same original model (Loeuille and Loreau, 2005) and all inspired by the Adaptive Dynamics framework, they can consistently vary regarding some of their assumptions, for instance regarding how the main evolving trait (body size or mass) is modeled: on a linear or a logarithmic scale (see Table 1 and below for an extended review). Given that these models can indeed typically result in the emergence of a trophic network, one can argue, on the one hand, that they capture the fundamental mechanisms responsible for food webs evolution and diversification. On the other hand, because of important variations between models, one can also argue that it is not clear which mechanisms are key or not, casting doubts on the generality of the models. For instance, results are very sensitive to the size of mutations considered: Allhoff et al. (2015) showed that the evolution of trophic network largely depends on the size of mutations, questionning the plausibiltiy of evolutionary branching as an important process underlying the evolution of trophic networks. It has also been shown that evolutionary branching critically depends on the choice of trade-off functions (de Mazancourt and Dieckmann, 2004). On the contrary, some parameters such as the amplitude of competition kernel or the density dependence on resources appear to have a negligible effect on diversification and the number of trophic levels (Brännström et al., 2011). In addition, it is still possible that the evolution of food webs observed in these models is due to strong and arbitrary assumptions shared among models (e.g. only one trait evolves or body size is arbitrarily lower bounded, see Table 1).

In this paper, we aim at better identifying the key mechanisms and assumptions that promote the emergence of trophic network in the class of eco-evolutionary models derived from Loeuille and Loreau (2005). We first review and thoroughly compare the different hypotheses made by these models. We especially show that some hypotheses are arbitrary and not justified from an ecological point of view. Second, we propose an unifying framework which includes all of the published models. Third, we show how relaxing strong assumptions made by previous models can give rise to degenerate trophic networks. We especially show that relaxing artificial bounds on evolving traits is critical and can dramatically change models’ predictions. Finally, we illustrate the sensitivity of the models by focusing on the assumptions made about how biomass is converted into new individuals. We argue that, given the lack of robustness of these models, their predictions should be taken with caution, especially if one aims at using these models for management of conversation purposes.

## 2 A review of food-web evolution models following Loeuille and Loreau (2005)

Eco-evolutionary models of food-web evolution derived from Loeuille and Loreau (2005) are inspired from the adaptive dynamics framework. They all share the same basic assumptions but show more or less important variations (see Table 1). These models are based on the modelling of the evolution of clonal species by introducing mutations into a population at ecological equilibrium. Ecological dynamics are given by a set of deterministic Lotka-Volterra equations, including resource consumption, predation, competition, birth and death. A resource is also considered, with its own dynamics (chemostat or logistic growth), which does not feed on any other species or resources. The main idea behind these models is that food webs structure is driven by traits of the species, especially individuals’ size or mass. Other traits are also considered, such as the prefered relative size of preys and specialization of predators. In general, higher dimensional trait spaces are known to promote branching (Ispolatov et al., 2015; Doebeli and Ispolatov, 2017). Allometries are also considered in all models, *i.e.* parameters, for instance individual death rate, are scaled with the size of individuals, following well-known empirical relationships between many traits (population density, longevity, reproduction rate, etc.) and size or biomass (Peters, 1983).

In all models, at least size (or biomass) is affected by mutations and evolves, assuming separation of timescales between ecological and evolutionary processes: mutations are introduced one after another when populations are at ecological equilibrium. The evolution of food webs is thus considered on long timescales. The effect of a mutation is drawn from a random distribution (Uniform or Gaussian, centered on the value of the parents or not). A mutation can be favored or not depending on the species already present in the environment. If it is favored, the mutant population invades and either replaces resident species or coexist with at least one other species. The analysis of the model is generally conducted thanks to numerical computations following a Piecewise-Deterministic Markov Process: population dynamics are deterministic and random mutations are introduced at random or fixed times.

All models assume that a single species is initially present in the environment, with the resources. Two phenomena can produce species diversification and the emergence of a food web. When only small mutations are considered, the food web gets enriched by a new species through evolutionary branching (*sensu* Adaptive Dynamics). When large mutations are possible, a new species may by chance appear in trait regions allowing invasion and coexistence with extant species. In all cases, models show that for a given set of parameters, the number of species can increase. Trophic interactions between species evolve since species i) can feed on other species, ii) have a prefered relative size of preys, iii) are more or less specialized on a prey size, iv) at least individuals’ size evolves. The results are hence generally presented as an interaction network, with the species as nodes, and trophic interaction between species as edges, with weights representing in- and out-flow of biomass. Finally, all models claim that for a set of reasonable parameter values, food-webs can emerge and evolve, and their structure can be close to trophic networks encountered in nature.

Despite all models are derived from the same seminal work by Loeuille and Loreau (2005), some of their assumptions can substantially vary. We now present in details how those models vary and why it matters. The differences are summarized in Table 1.

### Which traits evolve?

The number of evolving traits can differ, giving rise to a large variability of results. When only size evolves (Loeuille and Loreau, 2005; Brännström et al., 2011), resulting food webs are structured mostly linearly, and trophic levels are clearly defined. If size evolves with other traits as relative prey size preference or specialisation (Ingram et al., 2009; Allhoff and Drossel, 2013; Allhoff et al., 2015; Allhoff and Drossel, 2016; Bolchoun et al., 2017), food webs can show unsatisfying structures such as many species in a single trophic level, all feeding on the resource. Satisfying structures are then obtained under the assumption that mutations can be large. In the work of Ritterskamp et al. (2016), in addition to size, an abstract trait also evolves, which facilitate the emergence of food webs with different trophic levels and many different species. This shows that food web evolution largely depends on *a priori* assumptions about the evolving traits.

### Cannibalism, or not?

Cannibalism is excluded by some models, either because species are assumed to feed only on strictly smaller species (Loeuille and Loreau, 2005; Allhoff and Drossel, 2013), or because an abstract trait is supposed as structuring the food web and species with identical body masses / sizes can feed on species which do not share the same abstract trait (Ritterskamp et al., 2016). All other models allow cannibalism. *A priori* excluding cannibalism has several caveats. First, cannibalism is widespread in nature (Fox, 1975), and one can expect that models of food webs evolution reflect all features of observed trophic networks, including cannibalism. Second, excluding cannibalism artificially constraints the fate of mutants. For example, in the works of Loeuille and Loreau (2005) and Allhoff and Drossel (2013), individuals with a very small size difference can feed on and/or be eaten by each other while individuals with exactly the same trait can not. Why biomass flows are possible between very similar (but different) species and not for identical species is not clear. Furthermore, this introduces an additional selective pressure on neighbouring mutants: mutations close to their parent, which should be beneficial in cases without cannibalism (for instance by reducing direct competition), will usually be deleterious with cannibalism. Third, giving a justification of models excluding cannibalism based on individual-based models in the way of Costa et al. (2016) (see also Metz et al., 1996; Champagnat and Méléard, 2011; Champagnat et al., 2014; **?**) would require singular predation kernels which introduce artifical singularities in fitness landscapes. For instance, in the model by Ritterskamp et al. (2016), where an artificial trait is considered and cannibalism is excluded, considering only mutation along the abstract trait, the fitness landscape is always negative in the neighbourhood of the parent species, which implies that only mutants far away from the parents abstract trait can invade. This assumption artificially prevents the evolution of a continuum of species, with a continuous variation of abstract trait in the food webs, and thus artificially generates discrete species and discrete trophic interactions.

### Ordered predation

Some models (Loeuille and Loreau, 2005; Allhoff and Drossel, 2013) *a priori* assume an order for predation: species can only feed on smaller species. It necessarily excludes cannibalism and introduces artificial constraints (see before). Ordered predation also arbitrarily implies that in the case of species with very similar size, only the larger can feed on the smaller. Consequently, in the case of small mutations, it prevents trophic interaction changes: a prey can not become a predator. On the contrary, in the case of large mutations, a prey mutant can instantly become a predator if its size is larger than the size of the predator. This also introduces an asymetry in the fate of mutants: larger mutants are predators of their parent species whereas smaller mutants are preys of their parent species, even though both species are very similar for small mutations. Overall, assuming an ordered predation kernel imposes a structure to the food web and strongly constrains the evolution of trophic interactions.

### Mass/size or log mass/ log size?

Models either assume that food webs are structured with individuals size/mass or log size/log mass. The choice is important since, on the one hand, if the food web is structured linearly with absolute size or mass, it implies that trophic interactions and predation rates depend on the absolute size difference between predators and preys (Loeuille and Loreau, 2005; Allhoff and Drossel, 2013). If this predation distance is not subject to mutation and is then the same for all species, relatively to their masses, small species feed on much smaller species whereas large species feed on very similar species. Furthermore, for a given and fixed prey size preference, a small species can not feed on any other species because no species would be smaller, hence giving artifical constraints to the minimal possible size of species. In addition, models considering a structure with the absolute size/mass also assume that the resource has a size/mass equal to 0, which prevents species to be smaller than the resource, a constraint that can be considered artificial. On the other hand, assuming log size / log mass (as in Brännström et al., 2011; Allhoff et al., 2015; Ritterskamp et al., 2016; Allhoff and Drossel, 2016) allows to describe trophic interactions and predation rates depending on the relative size between preys and predators through a predation proportion rather than a predation distance. Models on the log-scale seem to be more realistic for food webs with fixed predation distance/proportion containing a large extent of (log)-masses / (log-) sizes. The same remark applies to models with evolving predation distance/proportion, in particular when the evolutionary speeds of both traits are different. Moreover, models on the log-scale assume that the resource has a log size/ log mass equals to 0 which allows species to become smaller than the resource if log-masses / log sizes are allowed to reach negative values. Overall, it appears that food webs models would rather be structured following log size/ log mass in order to introduce the least artificial constraints.

### Boundaries

Models make different implicit assumptions regarding the boundaries of possible values of evolving traits. In models assuming a food web structure following an absolute size/mass, size/mass is implicitly assumed larger than the resource as the resource size is 0, while in models assuming a structure in log size/log mass, the values could take any positive or negative values (although the extant models consider mutation distributions preventing species to evolve to negative values). Models considering other evolving traits, *i.e.* size preference for predation or predation specialization, also assume boundaries. The reason why such boundaries are assumed is not clear. It appears in some cases that if no boundaries constraints are assumed, the models generally give aberrant or trivial resulting trophic networks (see Allhoff and Drossel, 2013 where final food webs contain only one hyper-specialized species or a single trophic level; this is also mentioned by Allhoff and Drossel, 2016). If giving constrained boundaries to evolving traits is necessary to avoid unsatisfying results, the validity of such a theoretical framework can be questioned. On the contrary, one can expect that if such models capture the fundamental mechanisms underlying the evolution of food webs, then traits should evolve to realistic values on their own, without being artificially constrained.

### Mutation kernels

Mutation kernels are assumed either Uniform or Gaussian, generally centered around the parents value, but not necessarily: mutations can have effect proportional to the parental value (Loeuille and Loreau, 2005; Allhoff and Drossel, 2013) or independent of parental value for some traits (*e.g.* predation preference and predation specialization) but not for others (parents centered for size) (Allhoff et al., 2015; Allhoff and Drossel, 2016). Since models can make very different assumptions regarding their mutation kernels, it is difficult to compare them. For instance, assuming mutations not centred on the parental value (Allhoff et al., 2015) mimics migration from a regional pool of species and not mutations.

### Mutation size

All models, except the one of Brännström et al. (2011), assume that mutation effects are large. Some authors even show that if mutations are too small then diversification cannot occur (Allhoff et al., 2015). If mutation should have large effects otherwise food webs do not evolve accordingly to expectations, then evolutionary branching is certainly not an important phenomenon in the evolution of trophic networks. Indeed, evolutionary branching under the Adaptive Dynamics framework should occur for small mutations. If large mutations are necessary, other processes are involved. However, the model by Brännström et al. (2011) showed that small mutations can be sufficient to make emerge a satisfying trophic network. Hence, it is not clear why small mutations are sufficient under some conditions and not under others.

### Variations of other features

Models can also differ for other features: the dynamics of the resources, allometries or functional responses for predation rate. The resources can follow chemostat or logistic growth dynamics, with a feedback due to the decomposition of the dead individuals into resources or not; allometries can be introduced on one or several phenomena: birth, death, predation or competition rates; functional responses can follow Holling type I or II, or Beddington-DeAngelis. These varying assumptions may have strong influence on the evolution of food webs. Loeuille and Loreau (2006) have studied in particular the emerging allometries between species densities and bodymasses in the model by Loeuille and Loreau (2005). They concluded that the exponent of allometry is strongly influenced by predation parameters.

### Biomass conversion efficiency

Surprisingly, a single feature is common among all models: the *biomass conversion efficiency* (or conversion factor), that is the fraction of ingested biomass devoted to the production of biomass of newborns is supposed constant, independant of the mass / size of predators and ingested preys. This assumption is undoubtedly important because it implies that the mass converted by individuals when consuming preys is increasingly large when feeding on large preys, with no limit. In other words, there is no cost for predation: feeding on larger preys is not more costly than on smaller preys. This is in contradiction with empirical data which show that there is a trade-off for predators between eating small and large preys (Baras et al., 2014; Norin and Clark, 2017): there is an optimal prey size for which the conversion efficiency is the highest. This can be due to a trade-off between the low biomass given by small preys but lower costs to forage, handle and digest than for larger preys. It can also be due to the fact that predators can feed only on a part of a larger prey: the biomass converted from a large prey attains a maximum. This is also in contradiction with another common assumption of these models: if there exists a prefered size for preys, and if this preference can evolve, it should correspond to the best compromise between eating small or large preys. How the efficiency of biomass conversion affects the evolution of food webs has been ignored by extant models. We will show below that it is certainly a major mechanism underlying the evolution of trophic networks.

Overall, synthesizing the common and different assumptions of the models of food webs evolution derived from Loeuille and Loreau (2005) shows that the important mechanisms underlying the diversification and evolution of trophic networks are not clear. It is not clear whether there is a single major mechanisms that could give rise to satisfying structured networks, or whether combinations of different assumptions can give rise to similar results. This statement can cast doubt on the significance of such models for the study of food web evolution because models construction, analysis and exploration can be biased by *a priori* expectations of network structure and general features. This is not necessarily because some combination of assumptions give rise to satisfying networks structure that the underlying mechanisms are correct. At best, such models can give directions to what should be empirically verified: for instance, is it true that biomass conversion efficiency is constant? At worst, relaxing some of the assumptions made by the extant models, especially artificial constraints such as excluding cannibalism or imposing boundaries to evolving traits, could totally change the models’ results and give aberrant, trivial or unrealistic networks structure. This is what we will explore in the following. We first propose a general model, unifying the different extant models. Second, we use our model to explore the results given by extant models when relaxing some of their assumptions. Finally, we use our model to explore the importance of the conversion efficiency.

## 3 A unifying model

All the models reviewed in the previous section consist in specifying dynamics for species densities between mutation events. Between two mutations, assuming that the food web is composed of *n* species characterized by their average body mass *r*_*i*_ at maturity, we denote by *N*_*i*_ the population density of species *i* for all *i* = 1 *… n*. The resource density and biomass are indexed by *i* = 0. The basic dynamics for *N*_*i*_ takes the form

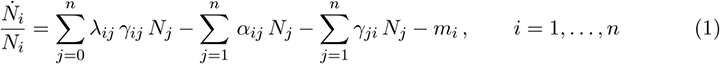

where *γ*_*ij*_ is the consumption rate of prey *j* by predator *i*; *m*_*i*_ is the mortality rate of species *i*; *α*_*ij*_ is the exogeneous direct competition rate between species *i* and *j*, which does not depend on resource and species consumption (sometimes called interference competition, Brännström et al., 2011); finally, *λ*_*ij*_ is the reproduction efficiency per capita of the consumer *i* and per capita of ingested biomass of species *j* (this quantity summarizes the biomass conversion and the reproduction; it can be interpreted as the fraction of an individual *i* produced by the consumption of an individual *j*); the reproduction efficiency *λ*_*ij*_ is related to the *biomass conversion efficiency*, denoted *ξ*_*ij*_ (see Section 2 and Section 5.1 for further definitions), by 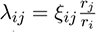.

The dynamics of the populations are coupled with the dynamics for a common resource density or biomass *N*_0_ which can be interpreted as organic or inorganic nutrient, with (bio-)mass *r*_0_. The dynamics of *N*_0_ varies among the references but it does not seem to significantly affect the behaviour of the model. In this work, in line with models by Brännström et al. (2011), we will consider the following logistic dynamics for the resource density:

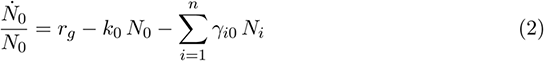

where *r*_*g*_ and *k*_0_ are the reproductive rate and the intraspecific competition rate of the resource population, respectively. In models by Loeuille and Loreau (2005); Allhoff and Drossel (2013); Ritterskamp et al. (2016), a chemostat equation is assumed for the dynamics of resources, with possible recycling of a fraction ν of biomass of dead individuals into resources:

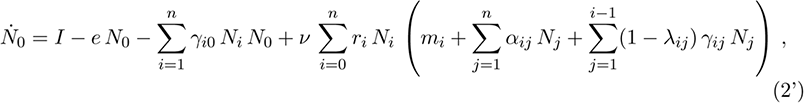

where *I* and *e*_0_*N*_0_ are the in- and out-flow, respectively.

In models (1) -(2) and (1) -(2’), each species is, *a priori*, allowed to predate any other species. The intensity of predation is governed by the function *γ*_*ij*_. For the ordered models, where predation of preys with larger body mass is forbidden (see Section 2), the predation rate *γ*_*ij*_ is 0 for species *i* larger than species *j*.

In order to make the link with the models of Loeuille and Loreau (2005) and Allhoff and Drossel (2013), which are expressed in term of biomass and with *r*_0_ = 0, we introduce the biomass of species *i* as *B*_*i*_ = *N*_*i*_ *r*_*i*_ and set *B*_0_ = *N*_0_. These quantities solve

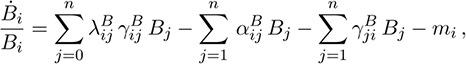

where

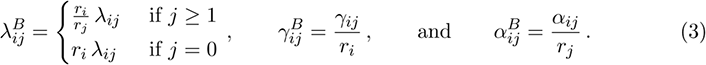

The ecological parameters *γ*_*ij*_, *λ*_*ij*_, *α*_*ij*_ and *m*_*i*_ are expressed in terms of individual parameters in the food web species including the body mass *r*_*i*_ or its normalized logarithm *z*_*i*_ = ln(*r*_*i*_*/r*_0_) (assuming *r*_0_ > 0), prefered distance (in terms of body mass) of predation *d*_*i*_ or its logarithm representation *µ*_*i*_ = ln(*r*_*i*_*/*(*r*_*i*_ − *d*_*i*_)) and a predation range parameter *s*_*i*_ on body mass scale or *σ*_*i*_ on logarithmic scale. Table 1 gives the different ecological parameters proposed in the cited references.

In all these references, mutations occur randomly in various ways, typically either with small probability at each time step or at regular time units. In all cases, they are assumed to occur on a long time scale, so that the system (1) and (2) or (2’) can reach a stationary state before next mutation. The species producing mutant is usually chosen with probability proportional to their density or biomass. The mutant trait is drawn according to the distributions given in Table 1 and introduced in the population at a given small density or biomass.

## 4 Revisiting extant models with relaxed constraints

### 4.1 A model with relaxed constraints

In order to investigate how relaxing the constraints reviewed in Section 2 might affect food web evolution, we will mainly focus on the model by Brännström et al. (2011) (see Table 1), which is structured on log scale, allows cannibalism and for which predation is unordered (species can feed on larger species). We consider that populations are characterized by their log-masses *z*_*i*_ and their predation preference *µ*_*i*_ (corresponding here to the predation log-distance) and assume without loss of generality that *r*_0_ = 1, so that *z*_*i*_ = log *r*_*i*_. The predation rate is given by

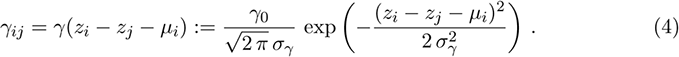

The favourite prey of species *i* has a log-mass *z*_*j*_ = *z*_*i*_ *-µ*_*i*_ or mass *r*_*j*_ when expressed as a fraction of its own body mass: 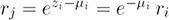. The shape of the predation rate is represented on the logarithm and linear scales on Figure 1.

**Figure 1:**
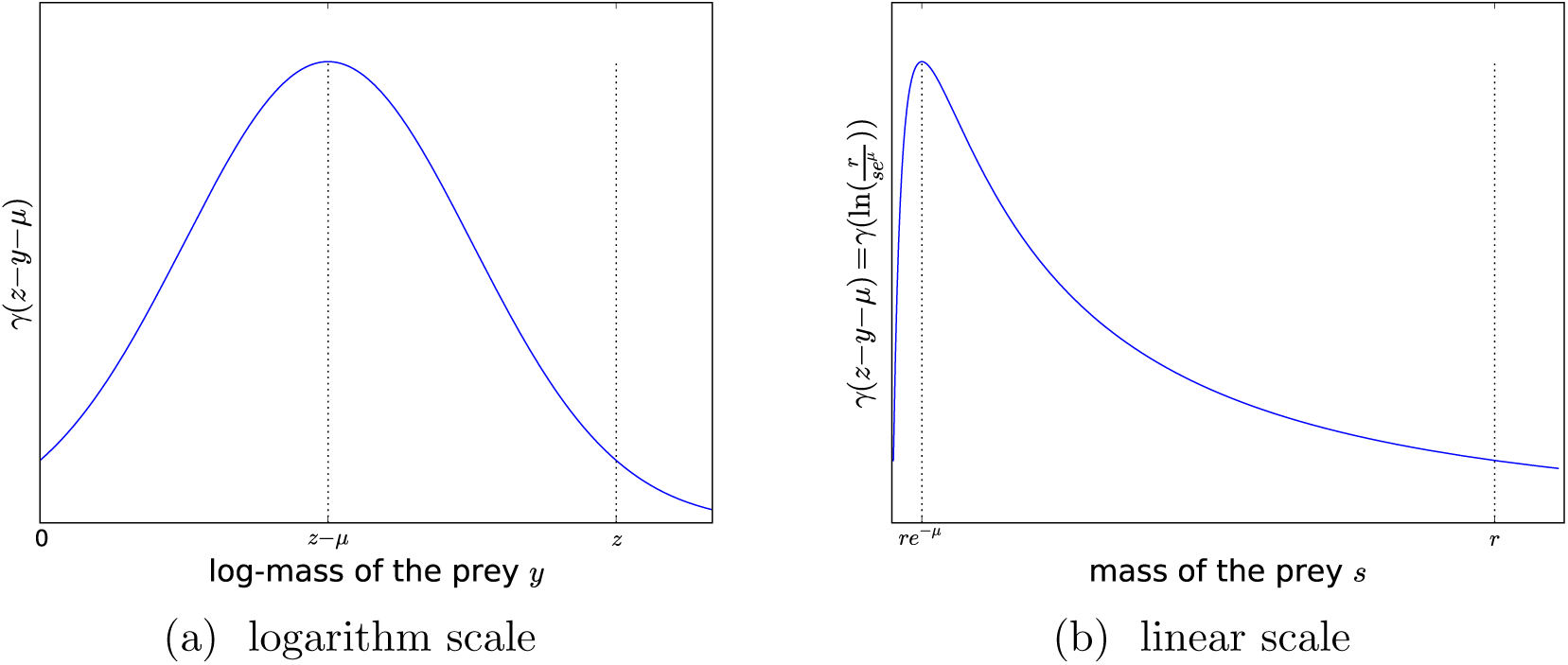
Predation rate (4) represented on the logarithm (left) and linear (right) scales.

**Figure 2:**
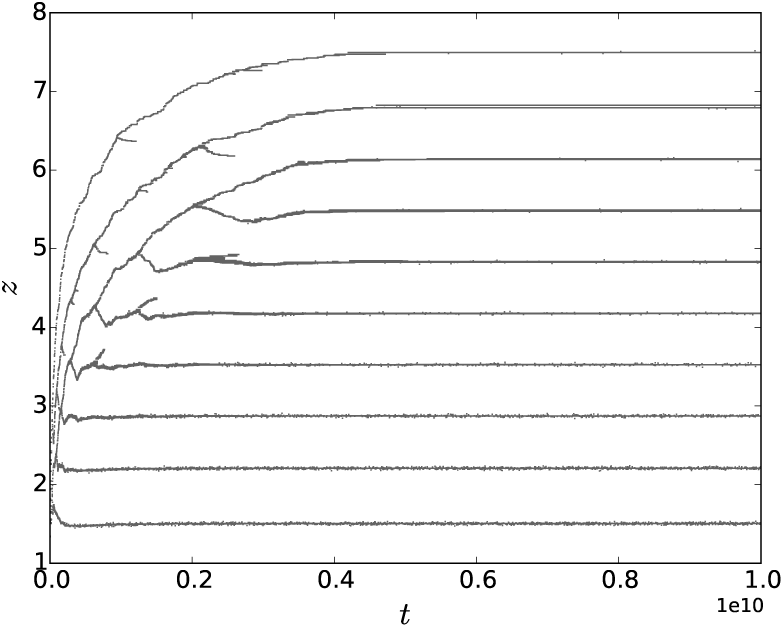
Evolution of the log-mass in the model by Brännström et al. (2011) with a single initial species with log-bodymass *z* = 1.2, variance of the mutation distribution *σ*_*z*_ = 0.01, predation preference *µ* = 3 and parameters of Table 2.

The competition parameter *α*_*ij*_ has the following Gaussian shape

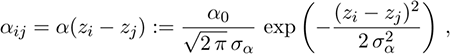

where *σ*_*α*_ is the competition range on the logarithmic scale.

The mortality rate takes into account allometry such as

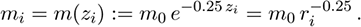

For reasons that we will explain in Section 5, we allow dependency on log-body mass of the conversion efficiency parameter (denoted by *λ*_0_ for extant models in Table 1) such that

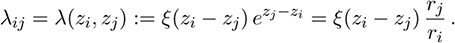

On the contrary to Brännström et al. (2011), both traits *z* and *µ* can evolve. We will see below that it yields unrealistic food web structures. In Section 5, we propose a solution to this problem, which does not require to let the predation range *σ*_*γ*_ evolve, contrary to what was claimed by Allhoff and Drossel (2013).

#### Methods^1^

The details of the computational methods slightly vary among references. Here we applied the following scheme.

Between mutation events, we compute the solution of Eq. (1)-(2) using the `odeint` python solver. Mutations are assumed to occur each *t*_*m*_ time units. This gives similar, yet faster, results to alternative schemes where mutations occur with a small probability at each time steps. The species (*z, µ*) producing a mutant is drawn proportionally to its density. The mutant (*z*’, *µ*’) is drawn such that *z*’ and *µ*’ are independant and Gaussian with means *z* and *µ* and variances 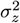 and 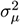 respectively. The mutant initial density is a small value *ε* and species are assumed to go extinct if their densities go below the same threshold *ε*. All the simulations in this paper are performed with *ε* = 0.0001. The density of the initial species and the initial resource concentration are the equilibrium of the system (1)-(2) with *n* = 1 given by

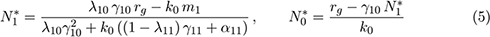

assuming that the fitness of the species 1 satisfies *λ*_10_ *γ*_10_ *r*_*g*_ − *k*_0_ *m*_1_ > 0, where *m*_1_, *λ*_10_, *λ*_11_, *γ*_10_ and *γ*_11_ are the death rate, production efficiencies and predation rates (for resource consumption and cannibalism) associated to the initial species.

The resulting food webs are represented such as an edge or a loop is drawn between predator *i* and prey *j* if predation of *j* by *i* is responsible for more than 10% (5% for dashed edges) of the reproduction of species *i*, i.e.

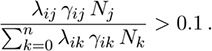

On Figures 3 and 5 to 9, a green edge means that the bigger species feeds on the smaller one and conversely for magenta edges. A loop is drawn if *i* = *j* and if more than 10% (5% for dashed loops) of the species reproduction is due to cannibalism.

**Figure 3:**
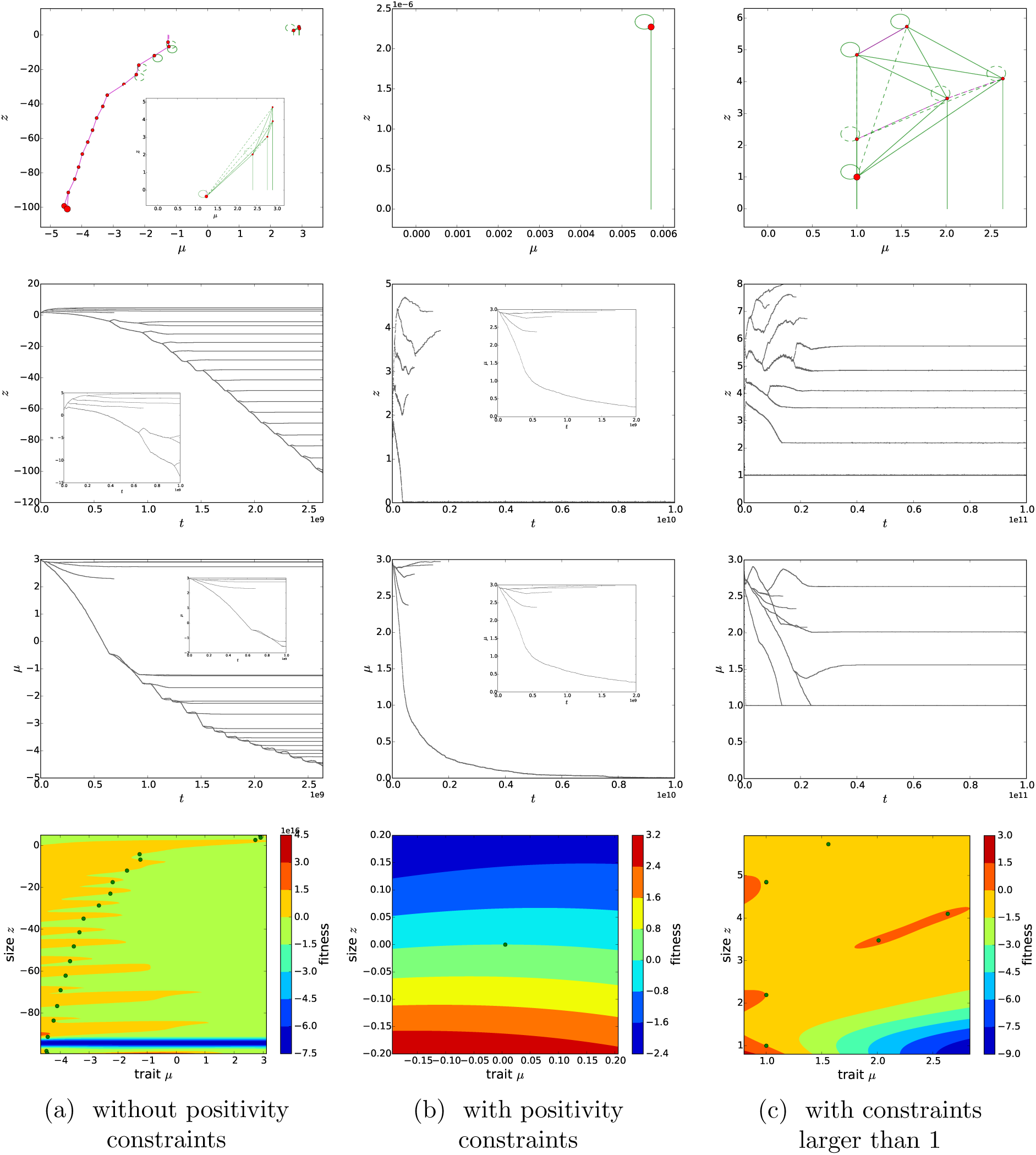
Food web evolution relaxing the constraint of fixed predation distance *µ*. From top to bottom: food web at final time of the simulation (*T* = 2.6 10^9^, overlay at intermediate time *t* = 4. 10^8^ (a); *T* = 1. 10^10^ (b); *T* = 1. 10^11^ (c)); evolution of the log-mass *z* (overlay: zoom on the begining of the simulation); evolution of the predation preference *µ* (overlay: zoom on the begining of the simulation); fitness lanscape at time *T* = 2.6 10^9^ (a); *T* = 1.0 10^10^ (b) and *T* = 1.0 10^11^ (c) letting *z* and *µ* evolving in the model of Brännström et al. (2011) (see Table 1 and Section 4.1) without (a), with (b) positivity constraints and with constraints larger than 1 (c) on evolving traits *z* and *µ* with parameters of Table 2, *σ*_*z*_ = 0.01 and *σ*_*µ*_ = 0.001.

### 4.2 Effect of relaxed constraints: unrealistic behaviours

In line with Brännström et al. (2011), with log-mass as the unique evolving trait, we obtain diversification by branching (see Figure 2) for identical parameters (given in Table 2), except for the range of competition *σ*_*α*_ which is a bit smaller in order to favour branching events (see Figure 9).

**Table 2:** Simulation parameters.

| Parameters | Values |
| --- | --- |
| $\sigma_\gamma$ | 1.5 |
| $\gamma_0$ | 10 |
| $\sigma_\alpha$ | 0.5 |
| $\alpha_0$ | 1 |
| $\lambda_0$ | 0.3 |
| $m_0$ | 0.1 |
| $r_g$ | 10 |
| $k_0$ | 0.01 |

Figure 3 shows three simulations where *z* and *µ* can both evolve (thus relaxing the hypothesis by Brännström et al., 2011, that *µ* is fixed), with much smaller mutations on *µ* than *z*. In Figure 3a, the food web initially evolves as expected: several branchings occur and the food web gets structured. However, the smallest species progressively evolves to smaller bodysize and predation preference until they both become negative. This means that this species feeds on a larger prey: the resource. After this, the richness of the positive part of the food web (i.e. the network composed of species with positive bodymasses and positive predation preferences) stops increasing and the negative part of the food web progressively diversifies, producing a linear food web with more and more negative traits. The stationary state of the food web is not reached at the end of the simulation, as the food web seems to evolve similarly endlessly. This behaviour is inconsistent with typical food webs that one may expect from such models.

As mentionned in Section 2, most of the previous models assume, explicitly or not, artificial constraints on the trait values. One may wonder if these constraints avoid such unexpected patterns. We ran simulations imposing positivity constraints on *z* and *µ* (by truncating mutation distributions below 0) and obtain Figure 3b. The behaviour of the food web is similar to the one of Figure 3a until the species with the smallest bodymass reaches zero. After this time, this body mass remains close to zero and the predation preference goes to zero. This produces a progressive loss of species ending finally with a single ‘resource-like species’, with body mass and predation preference close to zero. Again the behaviour is unrealistic and occurs for a wide range of parameters values. An interpreation for this behaviour is that the smallest species progressively adapts to the optimal consumption of resource. Since, in addition, the ‘resource-like species’ is subject to strong cannibalism, its density and the density of resource become too low for other species to survive. In Figure 3a, a small part of the positive food web remains because the bodymass of the smallest species becomes negative before being optimally adapted to the consumption of resources (*z* ≈ *µ*), hence a few amount of resources remain to sustain the survival of the positive part of the food web.

Replacing the artificial constraints on *z* and *µ* at 0 by a constraint at 1, we obtain Figure 3c. Contrarily to what Figure 3b shows, the food web has a non-trivial structure where several species progressively evolve to the constraint: three species have trait *µ* = 1 and one of them has also a log-mass *z* = 1. Due to the large values of production efficiency *λ* when predating large preys, we can observe in Table 3 that, although the proportion of predation due to cannibalism is small for most species in Figure 3c, the proportion of reproduction due to cannibalism is much higher (for example, for the species with traits *µ* ≈ 1.6 and *z* ≈ 5.7, the cannibalism represents only 0.9% of the global predation rate but 13% of the total species reproduction). Therefore, it is important to plot links between species based on criteria on reproduction rates rather than predation rates.

**Table 3:** Percentage of the total predation (top) and the total reproduction (bottom) of species by predation of other species in the food web at final time of the simulation *T* = 5. 10^10^ (see the top of Figure 3c) letting *z* and *µ* evolve in the model of Brännström et al. (2011) (see Table 1 and Section 4.1) with constraints larger than 1 on evolving traits *z* and *µ* with parameters of Table 2, *σ*_*z*_ = 0.01 and *σ*_*µ*_ = 0.001.

| Predators \ Preys | Resource | (2.6, 4.1) | (1.6, 5.7) | (1, 4.8) | (2, 3.5) | (1, 2.2) | (1, 1) |
| --- | --- | --- | --- | --- | --- | --- | --- |
| (2.6, 4.1) | 34.4% | 0.2% | 0.0% | 0.7% | 0.6% | 2.5% | 62.1% |
| (1.6, 5.7) | 9.1% | 9.2% | 0.9% | 6.1% | 10.6% | 9.3% | 54.8% |
| (1, 4.8) | 11.7% | 6.4% | 0.5% | 3.8% | 8.2% | 8.7% | 60.7% |
| (2, 3.5) | 34.4% | 0.2% | 0.0% | 0.07% | 0.6% | 2.5% | 62.1% |
| (1, 2.2) | 37.4% | 0.2% | 0.0% | 0.04% | 0.4% | 2.1% | 59.9% |
| (1, 1) | 51.0% | 0.03% | 0.0% | 0.0% | 0.09% | 0.9% | 48.0% |

| Predators \ Preys | Resource | (2.6, 4.1) | (1.6, 5.7) | (1, 4.8) | (2, 3.5) | (1, 2.2) | (1, 1) |
| --- | --- | --- | --- | --- | --- | --- | --- |
| (2.6, 4.1) | 12.7% | 5.5% | 0.4% | 3.1% | 7.3% | 8.3% | 62.6% |
| (1.6, 5.7) | 0.4% | 25.2% | 13.0% | 35.1% | 15.6% | 3.8% | 6.8% |
| (1, 4.8) | 0.7% | 25.1% | 10.2% | 31.3% | 17.0% | 5.0% | 10.6% |
| (2, 3.5) | 12.7% | 5.5% | 0.4% | 3.1% | 7.3% | 8.3% | 62.6% |
| (1, 2.2) | 15.0% | 4.0% | 0.2% | 2.1% | 5.7% | 7.6% | 65.5% |
| (1, 1) | 26.2% | 0.8% | 0.02% | 0.3% | 1.6% | 4.1% | 67.1% |

(b) Percentage of the total reproduction of predator species $(\mu_i, z_i)$ by predation on the resource or the preys species $(\mu_j, z_j)$ defined as $\frac{\lambda_{ij} \gamma_{ij} N_j}{\sum_{k=0}^n \lambda_{ik} \gamma_{ik} N_k}$ .

We emphasize that the artificial constraints that we put in the model in Figures 3b-3c play a key role in the food web evolution since some species reach the boundary of this constraint. Moreover, the sign of the invasion fitness (bottom of Figure 3) in the neighbourhood of these species shows that crossing these boundaries should be the natural evolution of the model.

Our results thus suggest that Loeuille and Loreau (2005) and Allhoff and Drossel (2013) obtained non-trivial food webs only because of artificial and hidden constraints due to the fact that *r*_0_ = 0 and *r*_*i*_ is positive for all *i* ≥ 1. In Allhoff et al. (2015) and Allhoff and Drossel (2016) the constraints on *µ* (and *σ*_*γ*_) are due to the mutation kernels which restrict them to an interval far from zero (see Table 1). They actually justify this choice because otherwise *µ* (and *σ*_*γ*_) can evolve to arbitrarily small values (see also Allhoff and Drossel, 2013).

Note that other unrealistic patterns were also observed in Allhoff and Drossel (2013) in the case where only *z* and *µ* evolve: the food web becomes composed mainly of a very large number of species on the first trophic level (with *z* ≈ *µ*) with a very wide range of bodysizes, and a few species on the second trophic level. We do not observe this kind of phenomenon, neither in the extension of the model of Brännström et al. (2011) proposed in this section nor in the one studied in the next section.

## 5 Illustrating food-webs sensitivity to hypotheses: the importance of biomass conversion efficiency

As shown in the previous section, the behaviour of the model is very sensitive to the parameters and the artificial constraints, may they be explicit or not. In some cases, the model’s behaviour is unrealistic and unexpected. The goal of this section is to investigate the influence of the biomass conversion efficiency — the only unchanged parameter among models in previous references — and the mutation kernel.

The shape of the biomass conversion efficiency is discussed in Section 5.1. A numerical study is performed in Section 5.2: we observe several threshold effects. In order to understand these thresholds we perform an analysis of fitness in Section 5.3, focusing on particular food web structures.

### 5.1 Necessity of a trade-off on conversion efficiency

In all previous models, the parameter *λ*_0_ of Table 1 is assumed to be independent of the body masses of the predator *z* and the prey *y*. This means that, regardless of the body-mass of the prey and the predator, the biomass produced by reproduction is a fixed fraction of the ingested biomass. The conversion efficiency is then increasing when the prey mass / size increases with no limit (see right panel of Figure 4, black thick line). This neglects numbers of tradeoffs, e.g. in term of energy used for the predation or handling time (see Section 2). In particular if individuals feed on larger preys, the hunting cost per unit of biomass is likely to be larger than for smaller preys. Moreover, the total biomass of large preys may not be ingested by predators. Conversely, if preys are very small, a predator has to feed on a large number of preys and then the handling time (per unit of biomass) becomes critical. This suggests that *λ*_0_ should depend on *z* − *y* as a function *ξ*(*z* − *y*), accordingly to empirical results (Baras et al., 2014; Norin and Clark, 2017), where *ξ*(*z* − *y*) should converge to zero when *z* − *y* converges to ±∞.

**Figure 4:**
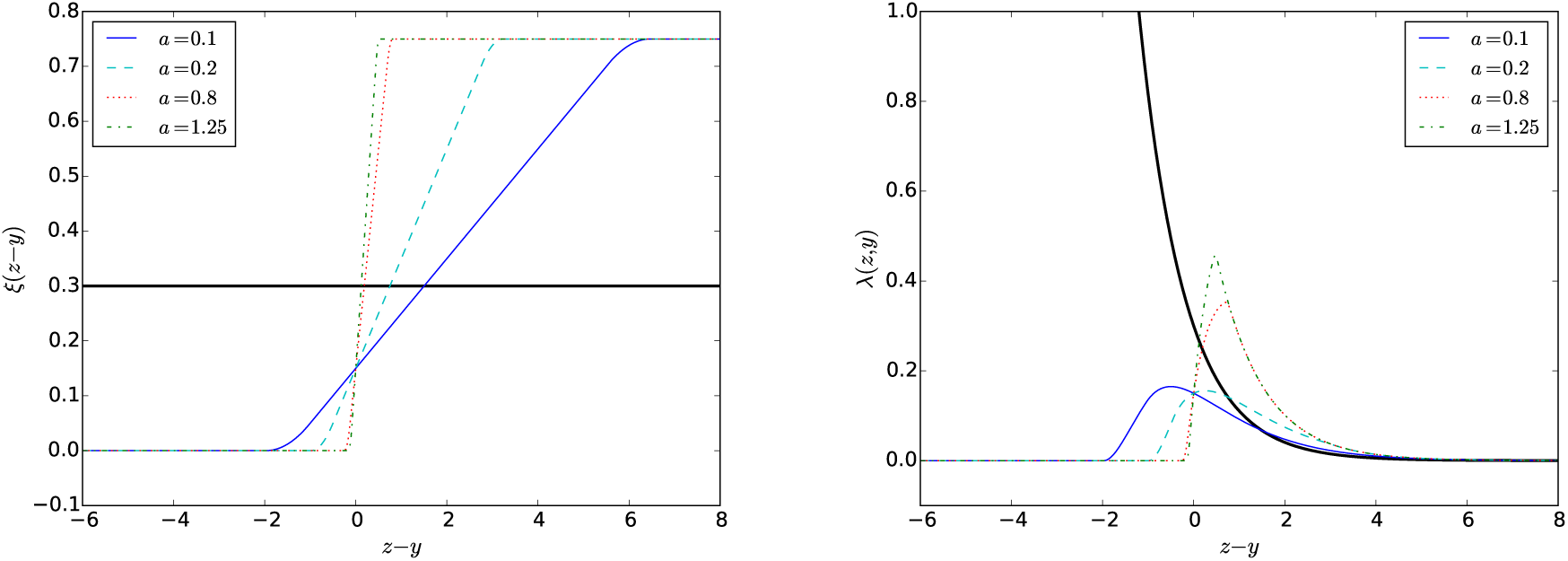
Possible shapes of the trade-off *ξ* (left) and *λ* (right) defined by Eq. (7) and (6) respectively, with *ξ*_max_ = 0.75 and *b* = 0.15 and for the model of Brännström et al. (2011), that is *a* = 0 and *b* = 0.3 (black thick line).

We assume the modified production efficiency function as following

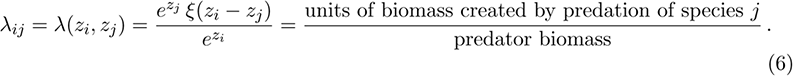

Recall that *λ*_*ij*_ represents the fraction of individuals of log-mass *z*_*i*_ produced by reproduction per unit of ingested individuals of log-mass *z*_*j*_. Since 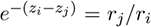, the term *ξ*(*z*_*i*_ − *z*_*j*_) represents the fraction of ingested biomass of species *j* devoted to reproduction of species *i*. We call *ξ*(*z*_*i*_ − *z*_*j*_) the *biomass conversion efficiency* of the ingested biomass into the biomass of newborns, through the predation of individuals of log-mass *z*_*j*_ by individuals of log-mass *z*_*i*_. Note that *ξ* is necessarily smaller than 1 to avoid the biomass creation *ex nihilo*, and was assumed constant in all the previous references.

### 5.2 Numerical study

Our numerical study shows that unsatisfying results obtained after relaxing strong assumptions (see Section 4) heavily depend on the behaviour of *ξ*(*z* − *y*) for small *z* − *y*. In our simulations, the range of predation preference in the food web is never large enough to be influenced by the decrease of *ξ*(*z* − *y*) for large *z* − *y*. A possible explanation is the limited amount of resources shared in the food web. For this reason, among hypotheses 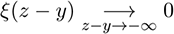 and 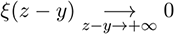, we will focus only on the first one. However, for models which behave as the one of Allhoff and Drossel (2013) (see last paragraph of Section 4.2), the second hypothesis is certainly to take into account.

Therefore, we focus on the following family of functions *ξ* parameterized by *a, b* > 0 and *ξ*_max_ ∈ (0, 1):

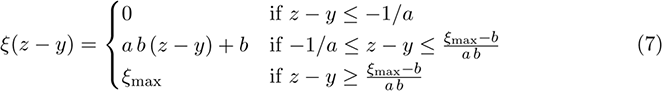

*ξ* is a linear function with slope *a b*, such that *ξ*(0) = *b* and truncated below 0 and above *ξ*_max_. We shall also assume that *b* < *ξ*_max_. This means that predation becomes harder in term of conversion efficiency in an interval of prey sizes containing the predator size. Examples of such functions are given in Figure 4.

To avoid problems of irregularity of fitness functions (see Section 5.3) we use regularized version of the previous curves (see Appendix A).

Figures 5 and 6 represent typical food web structures at the stationary state (except for four particular simulations, see legend) for several values of *a* and several speeds of evolution *σ*_*z*_ and *σ*_*µ*_ of the two traits.

**Figure 5:**
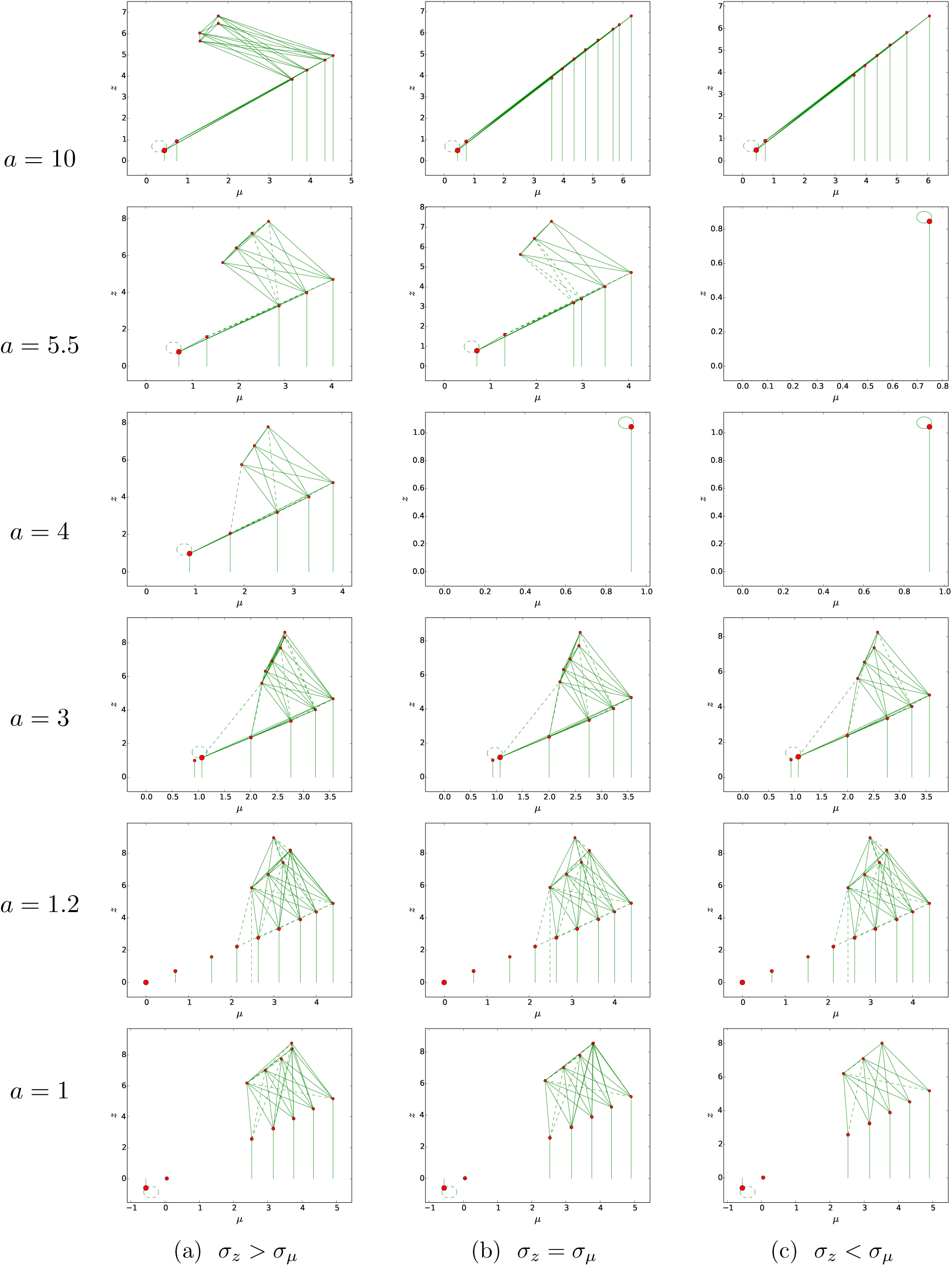
Food web at the stationary state for different values of *a* larger than 1 (except for *a* = 5.5 and *σ*_*z*_ *≥ σ*_*µ*_ and for *a* = 10 and *σ*_*z*_ > *σ*_*µ*_ which produce oscillations, see Figure 7) and for *σ*_*z*_ = 0.001 < *σ*_*µ*_ = 0.01 (a), *σ*_*z*_ = *σ*_*µ*_ = 0.01 (b) and *σ*_*z*_ = 0.01 > *σ*_*µ*_ = 0.001 (c). *b* = 0.15 and *ξ*_max_ = 0.75. Other parameters are given in Table 2.

**Figure 6:**
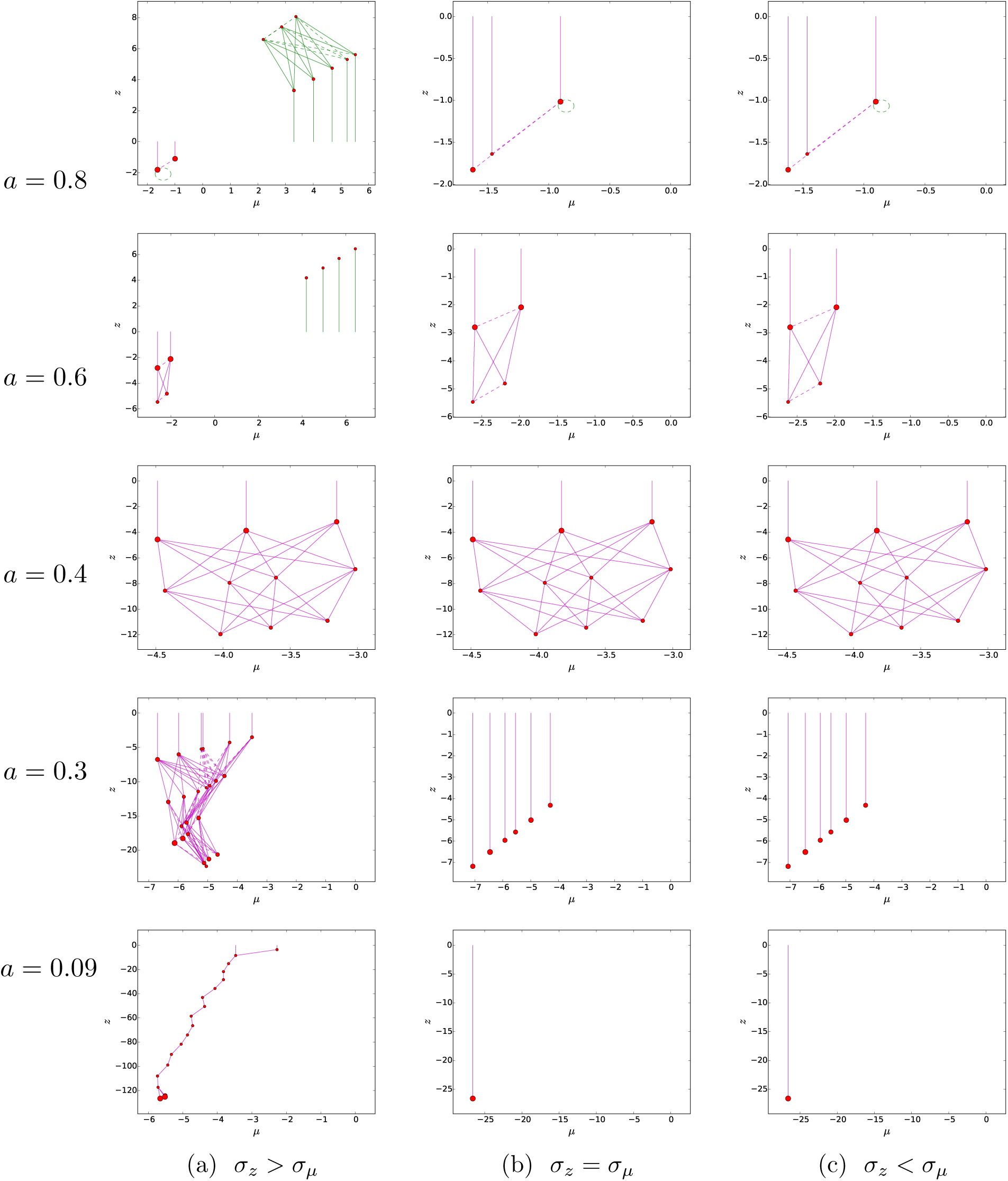
Food web at the stationary state (except for *σ*_*z*_ > *σ*_*µ*_, *a* = 0.03) for different values of *a* smaller than 1 and for *σ*_*z*_ = 0.001 < *σ*_*µ*_ = 0.01 (a), *σ*_*z*_ = *σ*_*µ*_ = 0.01 (b) and *σ*_*z*_ = 0.01 > *σ*_*µ*_ = 0.001 (c). *b* = 0.15 and *ξ*_max_ = 0.75. Other parameters are given in Table 2.

In Figures 5a and 6a, *σ*_*z*_ is bigger than *σ*_*µ*_, i.e. evolution is faster in the *z*-direction (although mutations are small in both directions). We observe for large values of *a* food webs with satisfying structures and several trophic levels. Since mutations are small, the community’s diversity grows due to successive evolutionary branching events. Progressively decreasing *a* (but still larger than 1.2), we observe the emergence of a new species with size very close to the resource and predation preference *µ* close to 0 (hence, subject to cannibalism). The emergence of such species seems to be a robust property of the model since it occurs for the three relative speeds of evolution of both traits (see others columns of Figure 5) and we experienced this phenomenon in our tests for a wide range of parameter values. We observe the convergence of the traits of the smallest species to (0, 0) when *a* approaches 1.2. This species does not reach negative values as long as *a* is larger than 1.2, although nothing in the model forces it to stay positive (contrary to the simulation of Figures 3b-3c). When *a* becomes a bit smaller than 1.2, this species has a negative log-mass and starts predating a prey (the resource) larger than itself. Meanwhile, the richness of the food web decreases progressively until *a* ≈ 0.6. For even smaller values of *a*, the part of the food web with positive biomass goes extinct and the richness of the negative part of the food web increases. In the limit *a* → 0, the conversion efficiency *ξ* converges to the constant function *b*. For small values of *a* (*a* = 0.09), we observe similar unrealistic behaviours as in Figure 3a. The fact that the food web becomes partly negative and the behaviour of the models becomes unrealistic at the threshold *a* ≈ 1 is explained in Section 5.3.1.

In Figures 5b and 6b (resp. Figures 5c and 6c) the simulations were run for equal ranges of mutations for both traits (resp. for larger mutations on trait *µ* than trait *z*). The behaviour observed for *a* > 1 (Figures 5b and 5c) is similar to Figures 5a except for intermediate values (*a* = 4 and *a* = 5.5) where the food web does not diversify: a single species consuming only the resource evolves towards an evolutionnary stable strategy. This is explained in Section 5.3.2.

A sensitivity on the mutation kernel is also observed when *a* < 1 (Figure 6). When *σ*_*z*_ = *σ*_*µ*_ or *σ*_*z*_ < *σ*_*µ*_, we still observe unrealistic food webs with negative bodymasses, but of different forms than in the case *σ*_*z*_ > *σ*_*µ*_. We may observe structured negative food webs as for *a* = 0.4, food webs with a single trophic level and a wide range of bodymasses as when *a* = 0.3 and *σ*_*z*_ ≤ *σ*_*µ*_, negative food chains as for *a* = 0.09 and *σ*_*z*_ > *σ*_*µ*_ or single species with very negative traits when *a* = 0.09 and *σ*_*z*_ ≤ *σ*_*µ*_.

The dynamics of some simulations are shown in Figure 7. They confirm that the food webs shown in Figures 5 and 6 are stationary, except for *a* = 5.5 and *σ*_*z*_ > *σ*_*µ*_, where periodic dynamics occur in the evolution of both traits *z* and *µ* (similar behaviour is observed for *a* = 10 and *σ*_*z*_ > *σ*_*µ*_ and for *a* = 5.5 and *σ*_*z*_ = *σ*_*µ*_) and for *a* = 0.09 and *σ*_*z*_ > *σ*_*µ*_, where the food web continues to evolve progressively to smaller negative body sizes as in Figure 3a.

**Figure 7:**
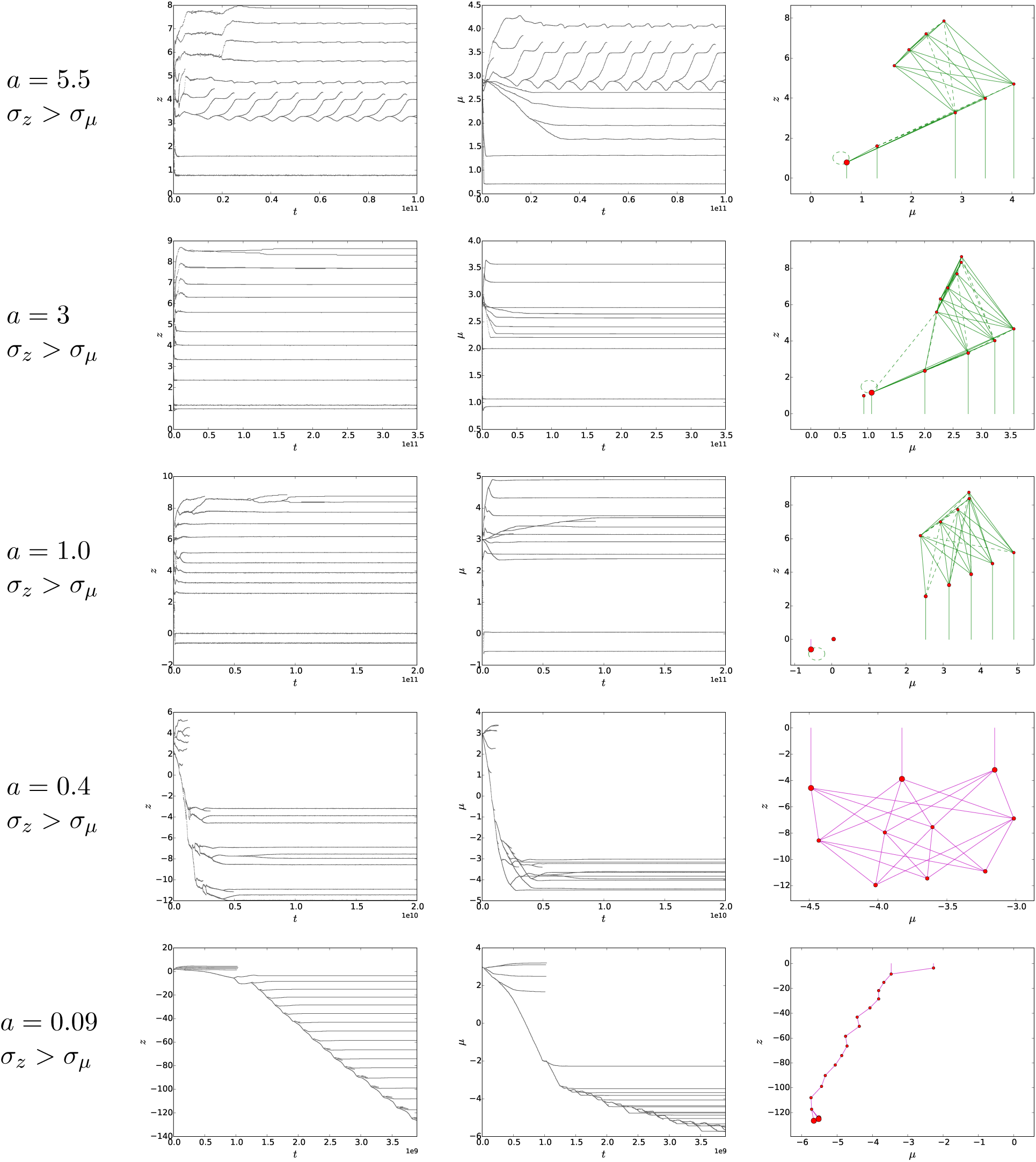
Evolution of the log-mass *z* (left), evolution of the predation preference *µ* (middle) and the final food web (right), for *ξ* defined by (7) with *ξ*_max_ = 0.75, *b* = 0.15, *σ*_*z*_ = 0.01 > *σ*_*µ*_ = 0.001 and *a* varying between 0.09 and 5.5.

We also see that, for values of *a* much smaller than 1, the food web first evolves to a realistic shape, similar to those observe for larger values of *a*, until the smallest body mass becomes too negative. After this, the positive part of the food web collapses and only the negative part remains.

Note that species close to (0, 0) suffer strong cannibalism. However, as the density of the resource is much higher than the density of the ‘resource-like species’, cannibalism does not contribute significantly to the species’ growth, which explains that loops are missing in some food webs of Figures 5 and 6.

We also ran simulations varying the parameter *b* of the conversion efficiency for a value of *a* avoiding unrealistic behaviour in Figure 5 (*a* = 1.2). The results in Figure (8) show a strong sensitivity of the stationary food web structure w.r.t this parameter. For all values of *b*, the initial food web dynamics shows progressive diversification as in Figure 5a, but when the smallest species become too close to 0, different behaviours are observed. For small values of *b*, the food web stabilizes in a realistic pattern. When *b* increases, the richness of the food web progressively decreases: species with intermediate bodymasses go extinct, until *b* = 0.4 where only three species with large bodymasses remain. For *b ≥* 0.45, only the ‘resource-like species’ survives and no further diversification occurs. This threshold effect will be explain in Section 5.3.3.

We also tested the sensitivity of the stationary food web to other parameters, in particular the variance *σ*_*α*_ of the competition kernel (see Figure 9). As observed in previous works (in particular by Brännström et al., 2011), this parameter has a strong influence on the richness of the food web: smaller *σ*_*α*_ promotes branching. Brännström et al. (2011) take *σ*_*α*_ = 0.6. For the numerical tests of Figure 5 and 6, we took a smaller value (*σ*_*α*_ = 0.5) to obtain richer food webs.

**Figure 8:**
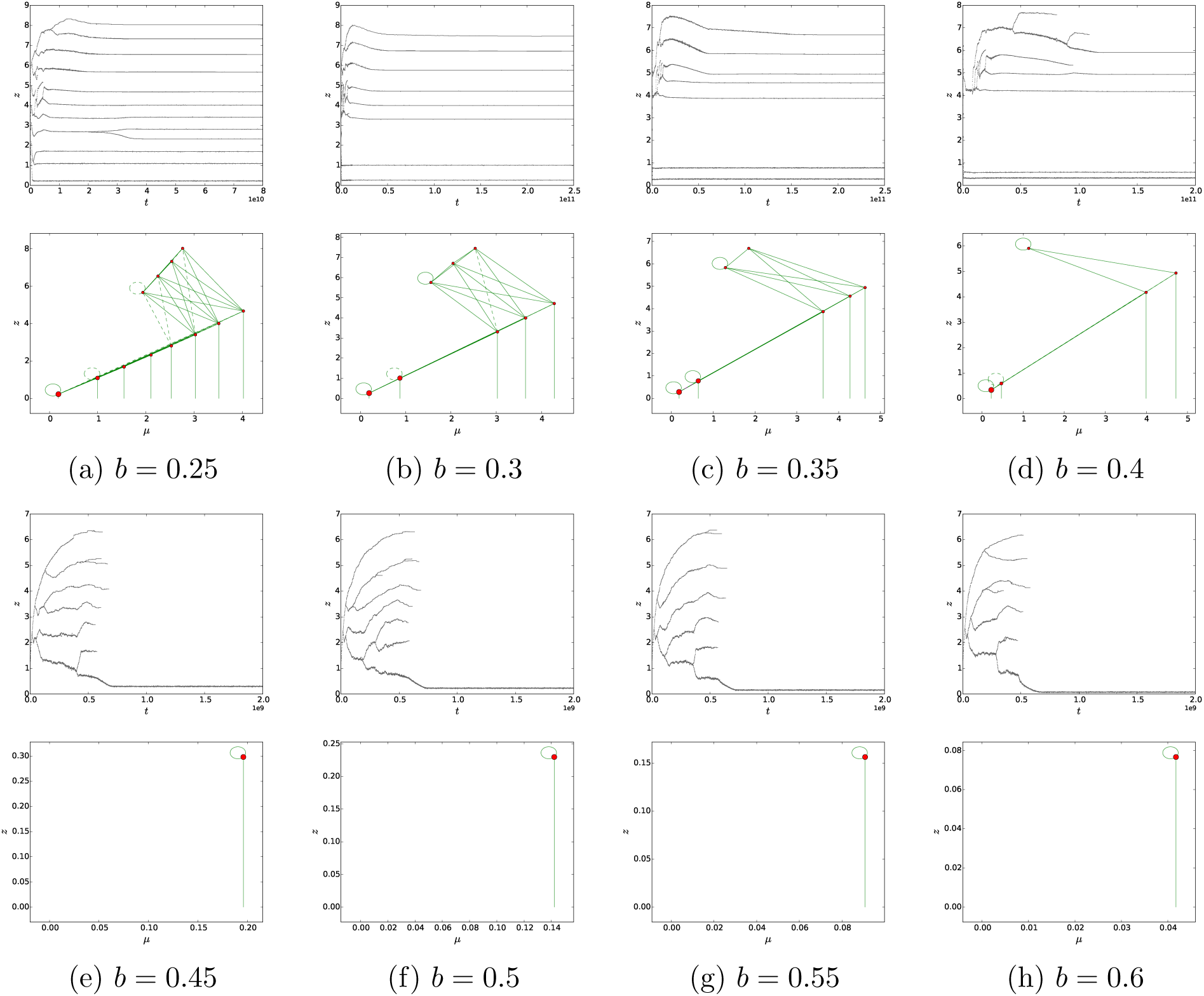
Evolution of the log-mass *z* (top) and food web at the stationary state (bottom) for *b* varying from 0.25 to 0.6 with *σ*_*z*_ = 0.01 > *σ*_*µ*_ = 0.001, *a* = 1.2 and *ξ*_max_ = 0.75. Other parameters are given in Table 2.

**Figure 9:**
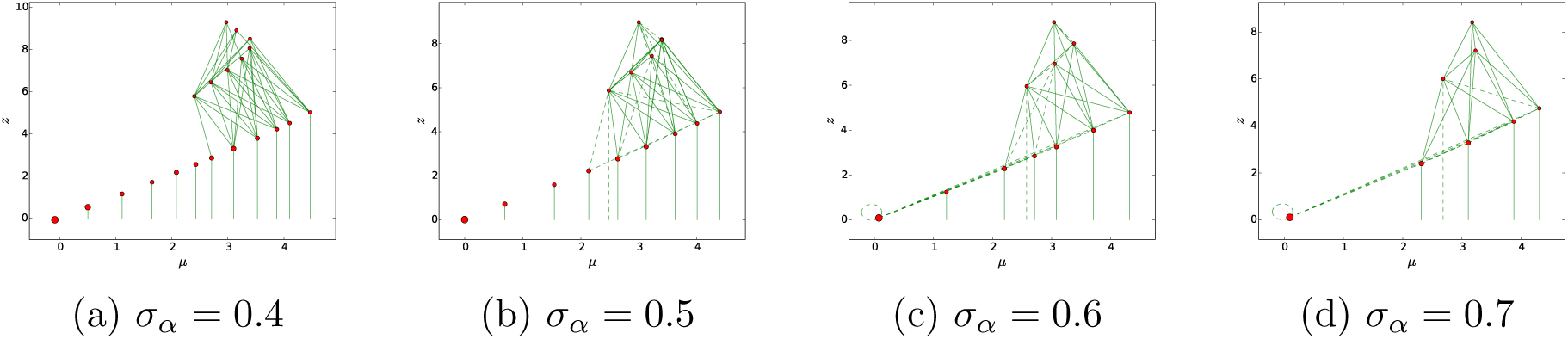
Food web at the stationary state for (a) *σ*_*α*_ = 0.4, (b) *σ*_*α*_ = 0.5, (c) *σ*_*α*_ = 0.6, (d) *σ*_*α*_ = 0.7 with *σ*_*z*_ = 0.01 > *σ*_*µ*_ = 0.001, *a* = 1.2, *b* = 0.15 and *ξ*_max_ = 0.75. Other parameters are given in Table 2.

### 5.3 Analysis of fitness

The invasion fitness of a mutant (*y, η*) in the food web (*z*_*i*_, *µ*_*i*_)_1≤*i*≤*n*_ is given by

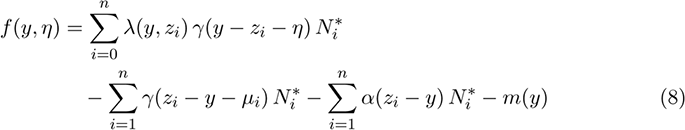

where 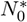 and 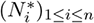 are respectively the resource concentration and the population densities at the stationary state of the food web, i.e. which nullify Eq. (1) and (2). The sign of the invasion fitness determines whether a species (*y, η*) can invade the food web (*z*_*i*_, *µ*_*i*_)_1≤*i*≤*n*_ or not. In particular we have the classical relation *f* (*z*_*i*_, *µ*_*i*_) = 0 for any *i* ∈ {1, *…, n*}.

#### 5.3.1 Threshold *a* ≈ 1

A key role in our analysis will be played by the derivative of the conversion efficiency:

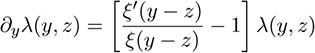

When *ξ* is constant (as in all previous references), we have *∂*_*y*_*λ*(*y, z*) = −*λ*(*y, z*) < 0. This means that there is a gain in reproduction when eating bigger species, or equivalently when the predator’s body mass decreases and the prey size remains constant. This is an important element to explain the global trend of evolution toward negative sizes. This also explains why the shape of the function *ξ*, and more precisely *ξ*’*/ξ*, may reverse this trend.

We can follow the adaptive dynamics’ paradigms to further analyse this effect: since we consider small mutations, the direction of evolution of a given species (*z, µ*) is governed by the fitness gradient ∇_(*y,η*)_*f* (*y, η*)|_(*y,η*)=(*z,µ*)_ (Metz et al., 1996; Geritz et al., 1998; Dieckmann and Law, 1996; Champagnat and Méléard, 2011; Champagnat et al., 2001),where

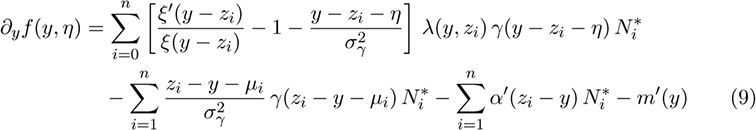

And

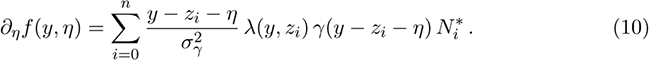

If *ξ* is constant, it follows from these expressions that a specialist species (i.e. a species whose growth is mainly due to a single prey) has a tendency to evolve toward smaller traits, provided that the speed of evolution of *µ* is fast enough compared to *z* and the density of its prey is large enough. Indeed, for such a species (*z, µ*), assuming that its major prey has size 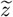 with density *Ñ*\* (potentially being the resource),

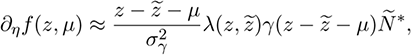

which makes the trait *µ* evolve to 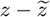 provided *µ* evolves fast enough, and then

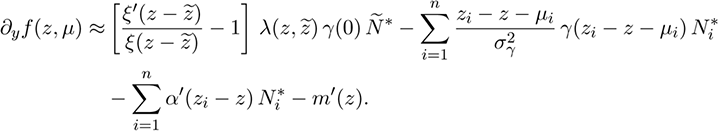

If we assume in addition that all species (*z*_*i*_, *µ*_*i*_) predating (*z, µ*) (i.e. such that *γ*(*z*_*i*_ − *z* − *µ*_*i*_) is not negligible) are also such that *µ*_*i*_ ≈ *z*_*i*_ − *z*, and that other species than (*z, µ*) have different enough sizes so that the main part of competition acting on (*z, µ*) is exerted by species (*z, µ*) itself, we obtain (using that *α*’(0) = 0)

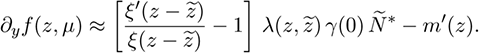

If *Ñ*\* is large enough, the last quantity has the same sign as 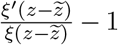, which is negative if *ξ* is constant. If 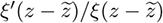 is larger than 1, this trend is reversed.

This argument applies in particular to cases with a ‘resource-like species’, i.e. a species with trait (*z, µ*) *≈* (0, 0). In the simulations of Figures 5, 6 and 7, the loss of richness of the food web and the appearance of unrealistic (negative) patterns seems closely related to the situations where a ‘resource-like species’ evolves to negative sizes. In these simulations, at the time where the resource-like species crosses (0, 0), the resource and resource-like species can be considered as a single species, which is relatively far from the rest of the food web, so that competition from other species is negligible and predation is exerted on this species only from specialist species. Therefore, letting 1 be the index of the resource-like species

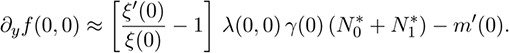

With *ξ* as in (7) and the parameters of Table 2, we obtain 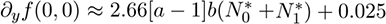. In all the simulations of Figures 5 and 6, the second term is negligible with respect to 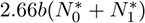 (for example, for *a* = 1.2, the latter is larger than 10). Hence the sign of *∂*_*y*_*f* (0, 0) is mainly given by the sign of *a* − 1, so that the resource-like species crosses (0, 0) if *a* ⪅ 1, leading to unrealistic food webs.

#### 5.3.2 First branching and sensitivity to the mutation kernel

We shall use the standard theory of adaptive dynamics to study the first branching event in the food web. We consider a single species (*z, µ*) and study the fitness of mutant traits (*y, η*). In this case, (8) becomes

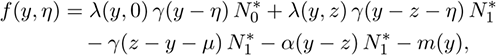

where we deduce from (5) that

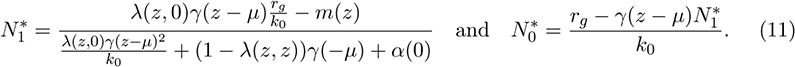

The possibility of evolutionary branching is linked to the existence of directions of local convexity of *f* (*y, η*) in the neighborhood of (*z, µ*) (Leimar, 2001). We plot in Figure 10 the signs of the fitness derivatives (see Eq. (14)-(17) in Appendix B) for *a* = 4 and *a* = 5 and for the three relative evolutionary speeds of both traits. At least in the beginning of the simulations, the initial species has a tendency to approach the curves *∂*_*y*_*f* = 0 and *∂*_*η*_*f* = 0, consistently with the classical theory of adaptive dynamics. Note that if *σ*_*z*_ = 10*σ*_*µ*_ (resp. *σ*_*z*_ = 0.1*σ*_*µ*_), evolution is much faster in the *z* direction (resp. *µ* direction) and the population first reaches the line *∂*_*y*_*f* = 0 (resp. *∂*_*η*_*f* = 0).

**Figure 10:**
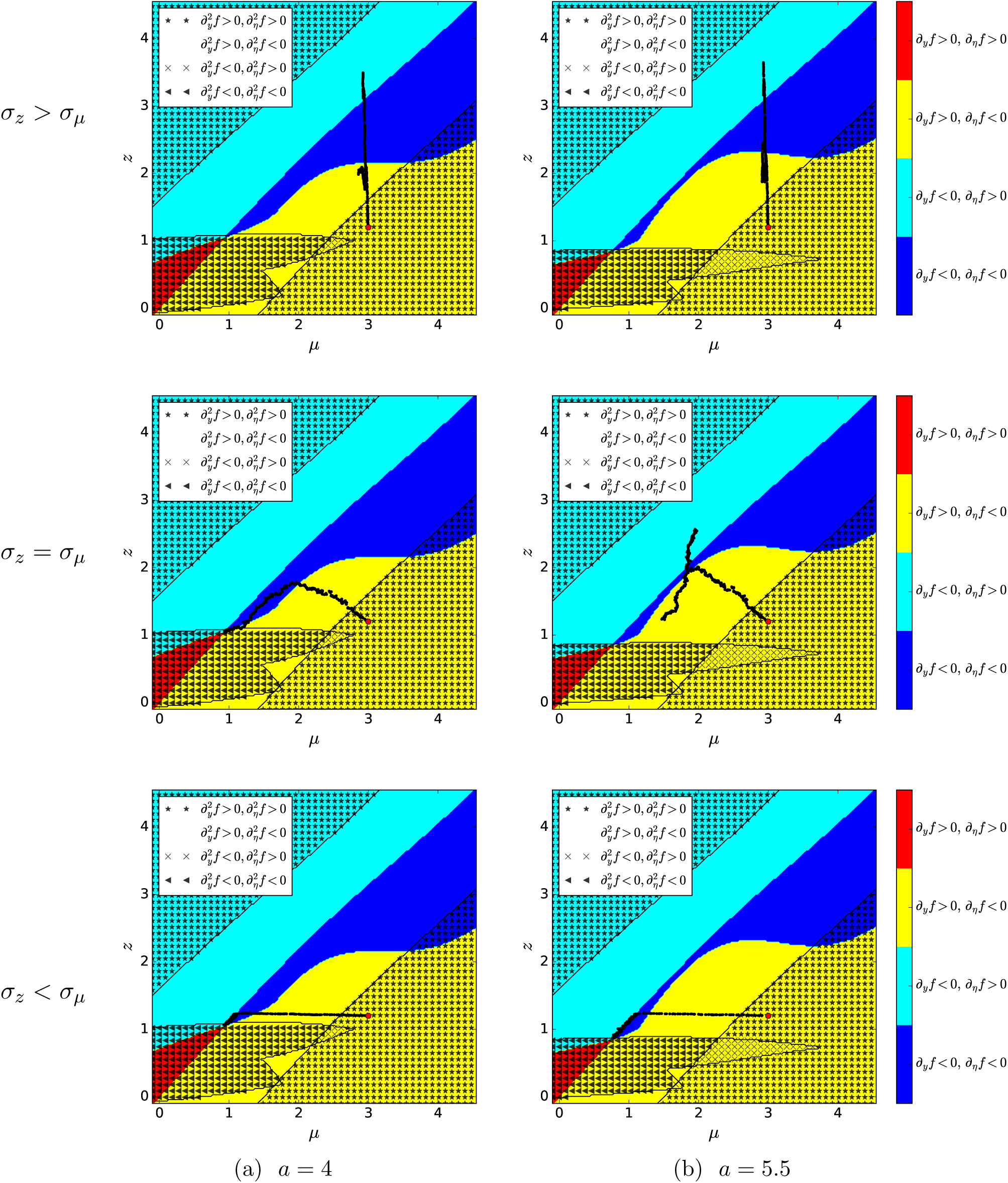
Sign of the first derivatives (14) and (15) and the second derivatives (16) and (17) of the invasion fitnesses for *a* = 4 (a) and *a* = 5.5 (b) and for 0.01 = *σ*_*z*_ > *σ*_*µ*_ = 0.001, *σ*_*z*_ = *σ*_*µ*_ = 0.01 and 0.001 = *σ*_*z*_ < *σ*_*µ*_ = 0.01 (from top to bottom). *b* = 0.15 and *ξ*_max_ = 0.75. The red dot corresponds to the starting point of the simulation and the black path to the initial evolution of the species, with possible branching.

We observe that 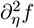 whenever *∂*_*η*_*f* = 0, so that the criterion of evolutionary branching is never satisfied when mutations only act on *µ* (see Appendix B). Therefore, evolutionary branching can only occur in the direction *z* of the trait space. Note also that, both for *a* = 4 and *a* = 5.5, the only evolutionary singularity in the region of traits we consider is located at the bottom left of the pictures, and is included in the region where both 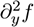 and 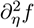 are negative. Hence, the evolutionary singularity is (locally) an evolutionary stable strategy and evolutionary branching cannot occur. Conversely, for smaller values of *a*, an evolutionary singularity appears in the region where 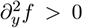 Hence this is a branching point, so that evolutionary branching occurs.

However, for *a* = 4 and *σ*_*z*_ > *σ*_*µ*_, and *a* = 5.5 and *σ*_*z*_ ≥ *σ*_*µ*_, we observe an evolutionary branching in Figure 5 which does not take place at the evolutionary singularity. It takes place along the line *∂*_*y*_*f* = 0 at points away from the curve *∂*_*η*_*f* = 0. This is explained by evolutionary branching along slow directional evolution, as described and analyzed by Ito and Dieckmann (2014). They claim that such evolutionary branching can occur along one direction of the trait space when the evolution in the orthogonal directions of the trait space is slow. The canonical equation of adaptive dynamics (Dieckmann and Law, 1996; Ben Arous et al., 2001) predicts that the speed of evolution in the *µ* direction of the trait space is proportional to the fitness gradient *∂*_*η*_*f* and the mutation variance 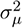. In cases where *σ*_*µ*_ = 0.1*σ*_*z*_, evolution in the *µ* direction is slow. When *σ*_*z*_ = *σ*_*µ*_, it is slow enough only for *a* = 5.5 because in this case the branching takes place at a point close to the line *∂*_*η*_*f* = 0, hence such that *∂*_*η*_*f* is close to zero.

#### 5.3.3 Sensitivity of the model with respect to *b* = *ξ*(0) and *ξ*_max_

In order to understand the collapse of the food web observed in Figure 8 for large values of *b* when the trait of the smallest species approaches (0, 0), we are going to consider the simpler situation of the extinction of the last species with large body-size *z*. Hence we assume that the food web is composed of the resource, a resource-like species for which we shall assume for simplicity that *z* = *µ* = 0 (i.e. resource consumption is optimal for this species), and a second species with traits *z* > 0 and *µ* > 0.

In this case, we shall use the competitive exclusion principle to decide whether the species (*z, µ*) is excluded by the resource like-species (0, 0). This will occur if the fitness *f* (*z, µ*) of the species (*z, µ*) is negative, where

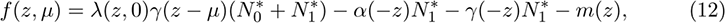

and 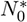 and 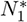 are the equilibrium densities of resource and resource-like species respectively, when the species with *z* > 0 is extinct. Hence for *ξ*(0) not too small (so that 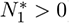)

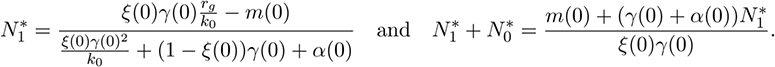

We then obtain

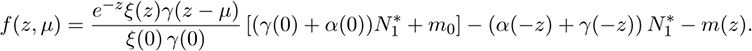

Then the species with *z* > 0 cannot survive if

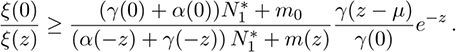

Observing that *ξ*(0) = *b* and assuming that *a* and *z* are large enough so that *ξ*(*z*) = *ξ*_max_ (and indeed the observed values for the body-size of the last ‘non-resource-like species’ in the food web is between 4 and 5 in simulations of Figure 8), we obtain, with the values of the parameters of Table 2,

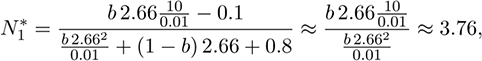

so that the species cannot survive if

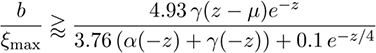

Therefore, we obtain a threshold effect for large values of *b/ξ*_max_, and we indeed observe in simulations values of the body size and the predation preference of the last surviving species with *z, µ* > 0 in the food web close to *z* = 4 and *µ* = 2.8, which gives, for *ξ*_max_ = 0.75 as in Figure 8, a threshold for *b* of approximately 0.41, above which we predict that the food web should collapse when the smallest species gets too close to trait (0, 0). This is consistent with the simulations of Figure 8. Biologically, when *b* increases, the resource-like species is more adapted for the resources consumption and for cannibalism. Then, for large *b*, the resource-like species is too competitive for other species to survive.

## 6 Discussion

In this paper, we first reviewed models of food webs evolution which followed the seminal work by Loeuille and Loreau (2005). These models are all based on the adaptative dynamics framework but vary in many of their assumptions, including arbitrary constraints on parameters and variables, or assumptions not compatible with the adaptative dynamics framework, e.g. the mutation size (see Table 1). We then propose a unifying model and we show 1) that under similar assumptions we recover the same results than previous models, and 2) that relaxing arbitrary constraints can lead to qualitatively different emerging food webs than the ones described in previous models. For instance, we find networks completely different than Brännström et al. (2011) when relaxing the assumption of fixed prefered predation distance, and than Allhoff et al. (2015); Allhoff and Drossel (2016) when relaxing the constraints on parameters range values. Finally, we show that a possible reason for the evolution of unrealistic food webs is that the biomass conversion efficiency is supposed constant in all models. Assuming on the contrary that the biomass conversion efficiency depends on the size difference between the prey and the predator can give satisfying foodwebs topologies: body sizes of the species remain larger than the basic resources size, and trophic levels emerge. We also finally show that trophic networks evolution strongly depends on the assumed form of the biomass conversion efficiency.

There are two manners to interpret our results. On the positive side, we can conclude that our results bring a lot in identifying the key mechanisms underlying the evolution of food webs. Indeed, we showed that simply considering a non-constant biomass conversion efficiency can solve most problems encountered with previous models. In addition, empirical data suggest that there is indeed an optimal size of prey size where predation is the most efficient (Baras et al., 2014; Norin and Clark, 2017). Our model is, in a sense, closer to data and observations and one might argue that such adaptive dynamics models really tell us how food webs emerge from eco-evolutionary processes. We can then go further into the analysis of our results and discuss about the importance of the biomass conversion efficiency and its form. We numerically explored a single family of function for the biomass conversion efficiency *ξ* (linear w.r.t. the size difference between prey and predator and truncated above and below fixed thresholds), but the fitness analysis shows that we can extend the observed results to more general functions. We exhibited the effect of two parameters on the expected trophic networks (Fig. 5, 6 and 8): *a*, the relative slope of *ξ* at 0 (i.e. *ξ*’(0)*/ξ*(0)) and *b*, the value of *ξ*(0) (Eq. 10, Fig. 4). The relative slope *a* and the parameter *b* both have threshold values at which networks evolve such as either all species have a size smaller than the resource (*a* ≃ 1), or the network is not stable and only one species remains (*b* ≃ 0.4 with our parameters values and for *a* = 1.2). For *a* smaller than 1, the evolution of the network towards smaller and smaller species is explained by the analysis of fitness (Section 5.3.1). When assuming that *ξ* is constant, as in all previous models (Tab. 1), the reproduction efficiency *λ*_*ij*_ of a species *i* feeding on species *j* increases exponentially when the size of the predator species *i* decreases (see Figure 4). This is because a given species converts more energy into reproduction as it feeds on large preys, especially on much larger preys than itself. Mutants decreasing size are thus necessarily favored and invade. Assuming a non-constant biomass conversion efficiency *ξ*, depending on the difference between the sizes of prey and predator species, our results show that: 1) *b* must be low enough to avoid the evolution to a network with a single resource-like species, and 2) *a* must be large enough to avoid too large benefits for species to feed on larger preys and to avoid the evolution to a network with species smaller than the resource. Our results would thus suggest that mechanisms underlying how energy is transfered from preys to predators, and how it is converted into predators biomass, are key for the evolution and stability of food webs.

However, one can interpret our results on a more negative side: our results can also suggest that such models are not robust enough to gain informative insights about food webs evolution. First because we show that relaxing hypotheses can lead to the emergence of qualitatively different, unstatisfying or unrealistic networks topologies. This shows that such models are very sensitive to assumptions. We showed that assuming a non-constant biomass conversion efficiency can solve problems but one can argue that such biomass efficiency can also evolve at the same time than body size and predation distance, and it might well be possible that completely different results would be obtained. Second, because one would expect that such models would be self-sufficient regarding certain general properties of trophic networks. In particular, if size effectively structures trophic networks with large species feeding on smaller ones, and if cannibalism is rare, this should emerge from the trait evolution and should not be due to arbitrary constraints. The model analyzed by Brännström et al. (2011) does not assume arbitrary constraints on the parameters range values, yet its results are not robust to the hypothesis of fixed prefered predation distance. Other models avoid cannibalism by *a priori* excluding it (Loeuille and Loreau, 2005; Ritterskamp et al., 2016), obtain hierarchically structured trophic networks by assuming that species can not feed on larger preys (Loeuille and Loreau, 2005), assume large mutations to observe network emergence (Allhoff and Drossel, 2013) or arbitrarily constrain parameters range values in order to obtain non-degenerated networks (Tab. 1). A general robustness analysis of such models appears necessary before generalizing their results, such as the one performed by Brännström et al. (2011). Finally we show, even after introducing appropriate forms of the conversion efficiency and in agreement with other authors, that results are very sensitive to mutation sizes, the number of evolving traits and the strength of interference competition, i.e. competition which is not due to resources and preys consumptions. In addition, the tradeoff we introduced on the conversion efficiency improves the behavior of the model where the predation distance *µ* evolves, but letting the niche width *σ*_*γ*_ evolve also requires to introduce delicate tradeoffs, as observed by Allhoff and Drossel (2013). Interference competition seems unreasonably necessary for the evolution of food webs: it is not clear whether there are good reasons why a non-trophic ecological process would be so important in food webs evolution, especially in models mostly based on competition for resources. Hence, our results altogether with results by previous authors can cast doubts on the explanatory and predictive power of such models.

One of the most important problem with the interpretation of such models is that it is difficult to avoid circular reasoning. In most previous papers, the authors claim that they are satisfied with the model’s results *because* they give realistic food webs structures. Some authors even justify arbitrary assumptions, such as a limited range for possible parameters values (e.g. Allhoff and Drossel, 2013), because otherwise results are not satisfying. We use ourselves such a circular reasoning to justify that assuming a non-monotonous biomass conversion efficiency *ξ* actually solves many problems encountered by the models. It is necessary to find ways to evaluate these models in a non-circular manner. A possibility would be to identify quantitative independent predictions that could be compared with food webs features. For instance, a limitation of the current models is that they *a priori* assume allometric relationships between size and parameters, generally as a power of 1*/*4 (Peters, 1983). However, one should observe that allometries measured in natural populations are *evolved* values and not fixed parameters. In models, including ours, allometry relationships are input parameters while they should be output parameters. We suggest that a possible way to evaluate such models would be to compare output allometric relationships to observed data as already done by Loeuille and Loreau (2006).

## Acknowledgements

This work was supported by the Chaire “Modélisation Mathématique et Biodiversité” of VEOLIA Environment, École Polytechnique, Muséum National d’Histoire Naturelle and Fondation X.

## A Regularization of *ξ*

To avoid problems of irregularity of fitness functions, we use in simulations a regularization of the function *ξ* defined by (7). The function *ξ* can be obtained as an affine transformation of the function *ξ*_0_ defined by

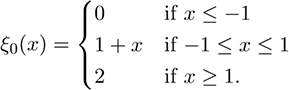

The regularization of *ξ* is obtained with the same affine transformation applied to the following regularization of *ξ*_0_.

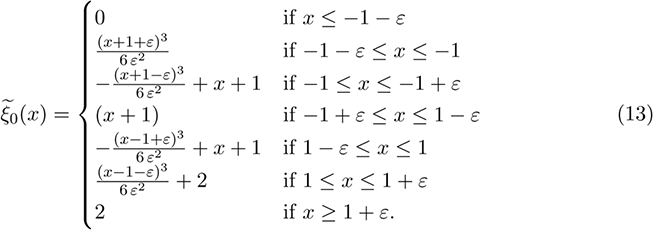

## B Second-order derivatives of the fitness

In the case where the food web contains a single species (*z, µ*) (see Section 5.3.2), the fitness gradient at the point (*z, µ*) is given by

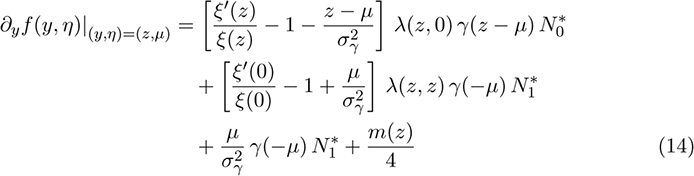

and

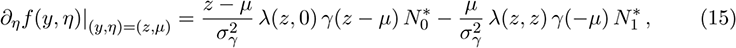

where 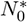 and 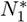 are given in (11).

The second derivatives of the fitness are given by

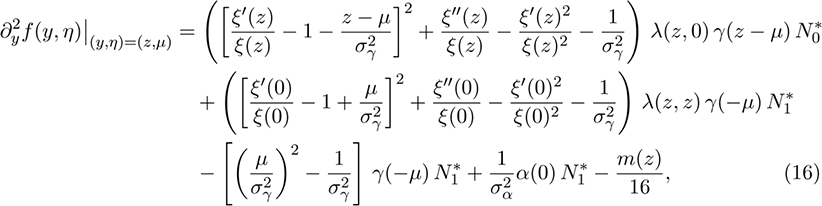

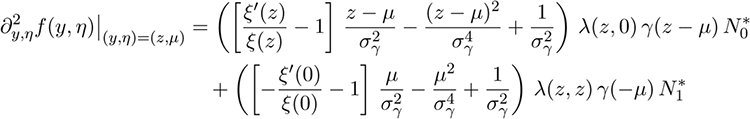

And

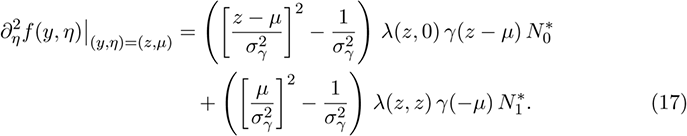

As observed in Figure 10, we deduce from these expressions that the curves 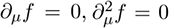 are close to the line *z* = *µ* and the pair of lines *z* = *µ ± σ*_*γ*_, respectively. This is due to the fact that, in the range of parameters we consider, the terms involving 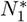 are negligible with respect to those involving 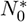. This implies that 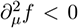 whenever *∂*_*µ*_*f* = 0, so that evolutionary branching cannot occur when mutations only act on *µ*.

## Footnotes

1 Simulations are run on the *babycluster* of the *Institut Élie Cartan de Lorraine* : http://babycluster.iecl.univ-lorraine.fr/

